# Phyletic distribution and diversification of the Phage Shock Protein stress response system in bacteria and archaea

**DOI:** 10.1101/2021.02.15.431232

**Authors:** Philipp F. Popp, Vadim M. Gumerov, Ekaterina P. Andrianova, Lisa Bewersdorf, Thorsten Mascher, Igor B. Zhulin, Diana Wolf

## Abstract

The bacterial cell envelope is an essential structure that protects the cell from environmental threats, while simultaneously serving as communication interface and diffusion barrier. Therefore, maintaining cell envelope integrity is of vital importance for all microorganisms. Not surprisingly, evolution has shaped conserved protection networks that connect stress perception, transmembrane signal transduction and mediation of cellular responses upon cell envelope stress. The phage shock protein (PSP) stress response is one of such conserved protection networks. Most of the knowledge about the Psp response comes from studies in the Gram-negative model bacterium, *Escherichia coli* where the Psp system consists of several well-defined protein components. Homologous systems were identified in representatives of Proteobacteria, Actinobacteria, and Firmicutes; however, the Psp system distribution in the microbial world remains largely unknown. By carrying out a large-scale, unbiased comparative genomics analysis, we found components of the Psp system in many bacterial and archaeal phyla and demonstrated that the PSP system deviates dramatically from the proteobacterial prototype. Two of its core proteins, PspA and PspC, have been integrated in various (often phylum-specifically) conserved protein networks during evolution. Based on protein sequence and gene neighborhood analyses of *pspA* and *pspC* homologs, we built a natural classification system of PSP networks in bacteria and archaea. We performed a comprehensive *in vivo* protein interaction screen for the PSP network newly identified in the Gram-positive model organism *Bacillus subtilis* and found a strong interconnected PSP response system, illustrating the validity of our approach. Our study highlights the diversity of PSP organization and function across many bacterial and archaeal phyla and will serve as foundation for future studies of this envelope stress response beyond model organisms.

## Introduction

The cell envelope is an essential, multilayered and complex structure, which physically separates bacterial cells from the environment. In their structural composition, Gram-positive and Gram-negative bacteria both share the cytoplasmic membrane and the cell wall. While the latter is much thicker in Gram-positive bacteria, the Gram-negative envelope additionally harbors an outer membrane (Silhavy et al. 2010). The cytoplasmic membrane is the functional barrier of the cell and fulfills crucial tasks, such as serving as a diffusion barrier, allowing the generation of the proton motive force and providing a platform for protein-protein interaction (Silhavy et al. 2010; Hurdle et al. 2011). Due to its essentiality, it is indispensable for prokaryotes to closely monitor and maintain their cell envelope integrity (Strahl and Errington 2017). This involves stimulus perception and signal transduction modules that comprise complex regulatory networks orchestrating a cell envelope stress response (CESR), which is activated when a cell is challenged with adverse conditions, such as envelope-perturbating antimicrobial compounds (Ulrich et al. 2005; Jordan et al. 2008).

One such system, the phage-shock-protein (PSP) response, has been studied in bacteria and one component of this system, PspA, has been identified in archaea and plants (Vothknecht et al. 2012). Initial studies in *Escherichia coli* revealed a strong induction of the PspA protein expression during phage infection accompanied by the production of the phage protein pIV, reassembling an outer-membrane pore forming secretin (Brissette et al. 1990). Subsequent studies on the PSP network identified various inducers including other secretins, elevated temperature or osmolarity, or interference with fatty acid biosynthesis (Bergler et al. 1994; Hardie et al. 1996; Kobayashi et al. 1998). In *E. coli*, PspA is encoded in the *pspABCDE* operon and expression levels are regulated by the PspF enhancer-binding protein via σ^54^ (Figure 1) (Brissette et al. 1991; Jovanovic et al. 1996). Under non-induced conditions, PspA forms a complex with PspF, thus silencing its own transcription (Figure 1) (Dworkin et al. 2000). Dependent on the stimulus perceived, PspB and PspC function as signaling units and initiate the disassembly of the PspA-PspF complex, thereby enabling activation of the system (Weiner et al. 1991; Kleerebezem et al. 1996; Flores-Kim and Darwin 2016). As a consequence, PspA proteins form a 36-meric donut-shaped oligomer that supports membrane integrity at the site of damage perception (Figure 1) (Engl et al. 2009). Interestingly, activation of the PSP system in *E. coli* by heat was shown to be PspB and PspC independent and solely required PspA (Weiner et al. 1991). Thus, PspA functions as i) a regulator, ii) a sensing unit, and iii) an effector protein, substantiating its key role within the PSP network. The biological function and physiological significance of the remaining PSPs, PspD and PspE, are still unclear (Adams et al. 2002; Flores-Kim and Darwin 2016). PspF also regulates the orphan *pspG* gene, which is the only other known PspF-target in *E. coli*. However, the role of PspG in the PSP response is not fully understood (Green and Darwin 2004; Flores-Kim and Darwin 2016). Deletions in the *psp* operon of *E. coli* show mild phenotypes, such as growth defects in late stationary phase (Weiner and Model 1994; Flores-Kim and Darwin 2016).

**Figure 1:**
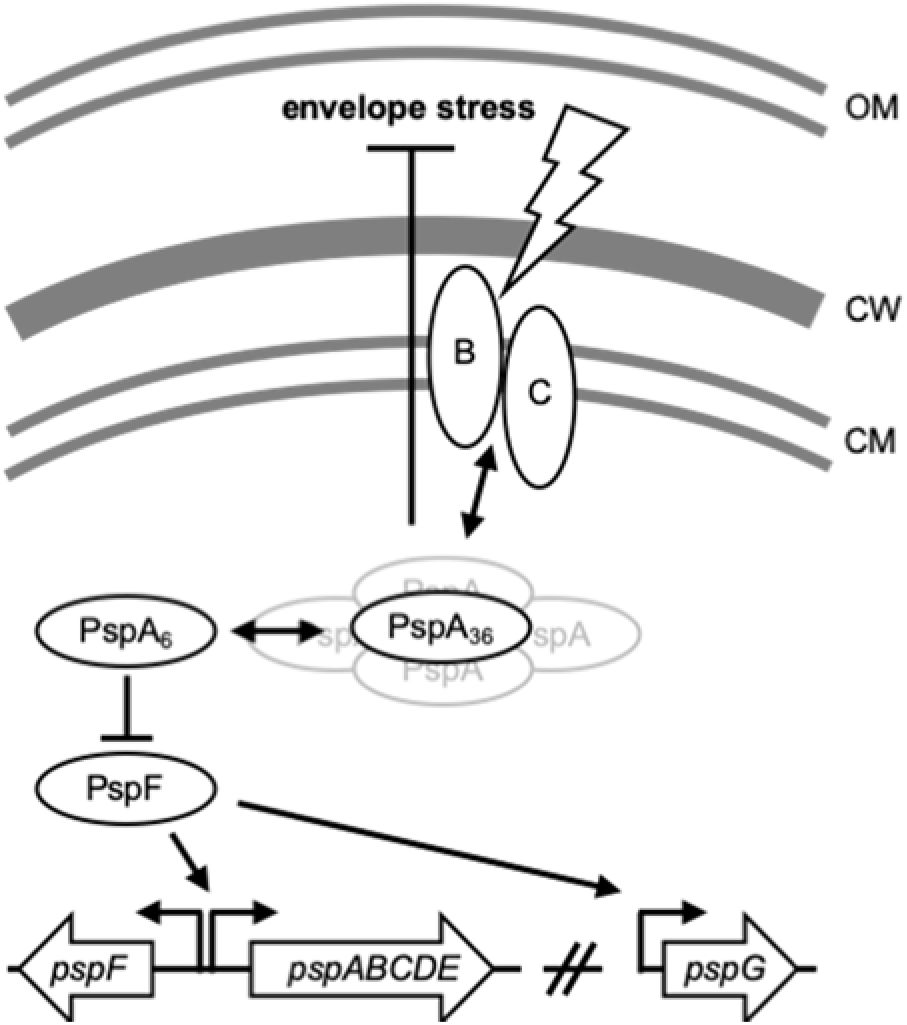
The phage shock protein response in *E. coli*. Upon stimulus perception mediated by the signal detectors PspB and PspC, PspA oligomerizes and presumably supports membrane integrity at site of damage perception. Transition of PspA polymerization state (either mediated by or independent of PspB/C) causes release of the transcriptional regulator PspF, enabling transcription of the *pspA-E* operon. In *E. coli*, the orphan *pspG* gene is the only other known target of PspF, however its biological function within the PSP response is still unknown.

In contrast, the PSP system in *Yersinia enterocolitica* is of importance for bacterial survival when the virulent Ysc type III secretion system is expressed during host infection (Darwin and Miller 2001). It has been shown that the deletion of *pspC* results in reduced virulence and growth defect (Darwin and Miller 1999; Darwin and Miller 2001). Detailed research on the PSP response also revealed the roles of the membrane proteins PspB and PspC as dually (positive and negative) acting regulators in *psp* operon expression in *Y. enterocolitica* (Yamaguchi and Darwin 2012). Surprisingly, PspA plays no essential role in terms of cell growth and bacterial survival during host infection, whereas PspBC are needed to protect the cells from secretin-induced death (Maxson and Darwin 2006; Gueguen et al. 2009). Furthermore, the genetic organization of the PSP locus in *Y. enterocolitica* differs from that in *E. coli*, e.g. PspC contains an N-terminal extension that is involved in *psp* gene expression regulation and is not present in *E. coli* and a PspE homolog is missing in *Y. enterocolitica* (Darwin and Miller 2001; Flores-Kim and Darwin 2016).

In the Gram-positive model organism *Bacillus subtilis*, the PspA homolog termed LiaH also forms oligomeric ring structures as a consequence of cell envelope stress (CES), thus substantiating the role of PspA-like proteins in supporting membrane integrity (Wolf et al. 2010; Wolf et al. 2012). In contrast to the regulation in *E. coli*, *B. subtilis liaH* is controlled by the two-component system (TCS) LiaRS, which strongly induces expression of the *liaIH* operon upon perceiving CES. In addition to LiaH, this operon encodes a membrane anchor protein LiaI, which facilitates LiaH recruitment to the cytoplasmic membrane (Jordan et al. 2008; Wolf et al. 2010). *B. subtilis* also encodes a second PspA-like protein and a PspC domain-containing protein in separate operons. Potential diversity of the Psp system is further supported by a recent study of the actinobacterium *Corynebacterium glutamicum*, where the PspC domain was found as the N-terminal input module of a histidine kinase, embedded in a three-component system responsive to CES (Kleine et al. 2017).

Previous genomics studies focused on analyzing the PSP system only in three bacterial phyla - Proteobacteria, Firmicutes, and Actinobacteria, with experimentally studied representatives (Huvet et al. 2011; Ravi et al. 2018), whereas the newest genome-based taxonomy defines more than 100 bacterial and archaeal phyla (Parks et al. 2018). Thus, our knowledge on diversity and distribution of the PSP system throughout prokaryotes is very limited. To fill this gap, we performed a large-scale genomic analysis of PSP networks in bacteria and archaea by analyzing over 22,000 genomes, representing all bacterial and archaeal phyla for which sufficient genomic data is available (Parks et al. 2018). First, we analyzed the distribution of PSP-specific domains throughout different phylogenetic ranks. We then performed an in-depth profiling of putative PSP networks encoded in each genome in the dataset. The PspA and PspC domains showed the highest diversity with respect to phyletic distribution and associations with other domains within a single protein. By analyzing the domain architectures and genetic neighborhoods of PspA and PspC, we identified new genomic organizations and provided context-specific knowledge enabling predictions of novel PSP network architectures and domain combinations. Using a broad bacterial two-hybrid screen we confirmed the results of our *in silico* analysis by experimentally dissecting the PSP network of *B. subtilis*, which consists of 14 known and predicted PSP proteins encoded in three separate genomic locations.

## Results and Discussion

### Genomic perspective on the phage shock proteins

The genomic sequence space of the microbial world is rapidly increasing with close to 150,000 bacterial and more than 2,000 archaeal genomes currently classified in the Genomic Taxonomy database (GTDB v86) (Parks et al. 2018). But the sheer size of this dataset does not necessarily reflect the phylogenetic diversity in nature, as the number of sequenced bacterial genomes is currently heavily biased towards three bacterial phyla: Proteobacteria, Firmicutes and Actinobacteriota, comprising more than two-thirds of the available genomic data (Parks et al. 2018). This leaves the remaining bacterial phyla highly underrepresented and demands an unbiased approach to tackle genomic data. We therefore first generated such dataset, containing approximately 22,000 genomes that represent 99 bacterial and 10 archaeal phyla (Figure 2A, and Material and Methods). This set of genomes is a balanced dataset compiled and used for classification by the GTDB (Parks et al. 2018). Next, we applied Hidden-Markov-Models (HMMs) of each phage shock protein (PSP) domain present in *E. coli* on all genomes to screen our dataset for the diversity of the PSP systems throughout bacteria and archaea. Then we analyzed the phylogenetic distribution of each domain from the known proteobacterial PSP network (Figure 2A). 83 bacterial and 7 archaeal phyla contain genomes encoding PSPs, but the abundance of PSP-positive genomes within these phyla varied substantially (Figure 2A, black/white circles). About 45% and 60% of the phyla contained genomes encoding the effector protein PspA and the signaling protein PspC, respectively. Their wide phylogenetic distribution was the first indication that the PspA and PspC domains represent the core of the PSP network architecture (Figure 1A). This was supported by the rapid descending number of phyla encoding other members of the PSP system, such as the signaling protein PspB, which was found only in 26% of the analyzed phyla.

**Figure 2:**
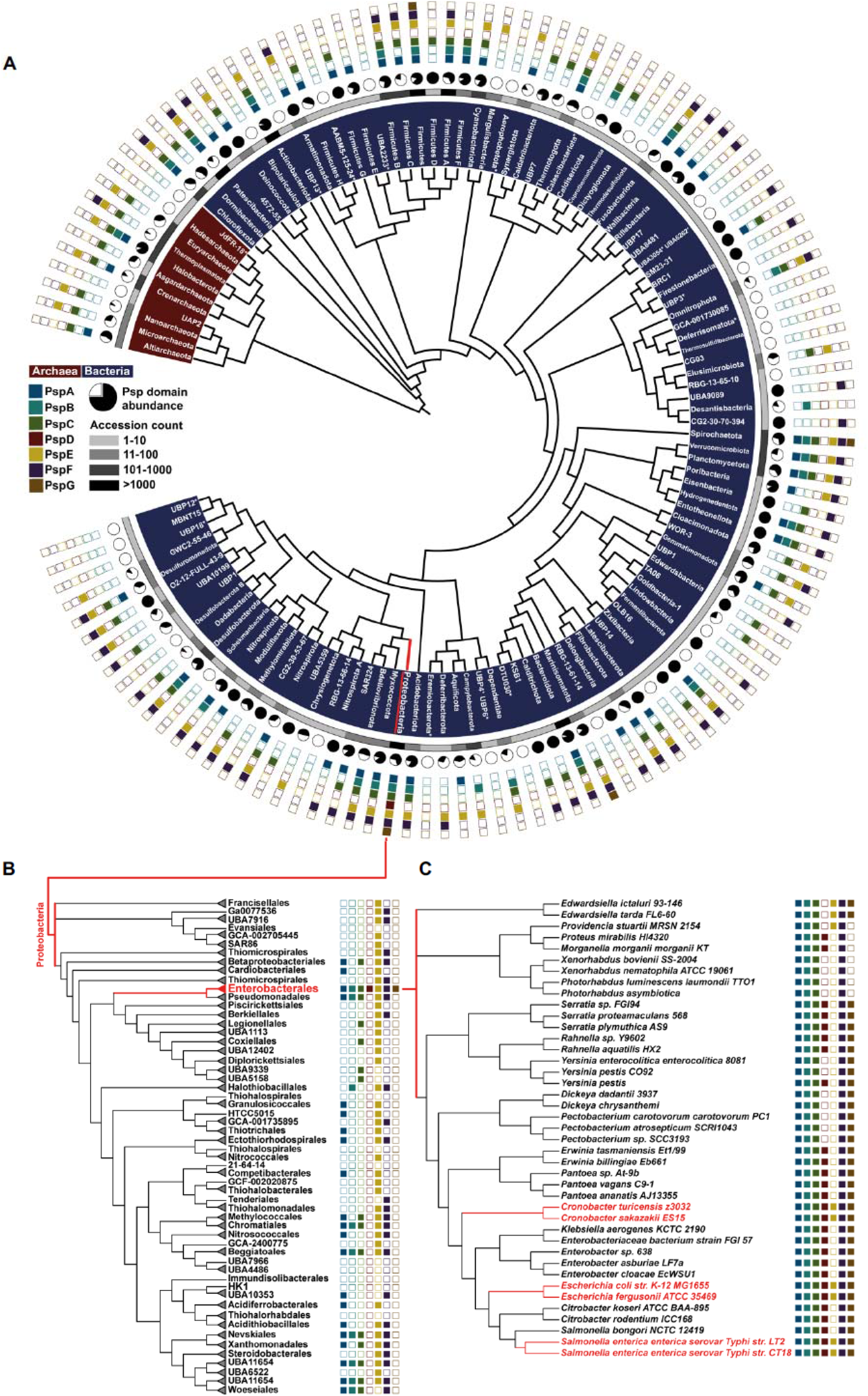
Phylogenetic diversification of PSPs in bacteria and archaea. **A** Phylogenetic representation of bacterial and archaeal phyla (phylogenetic tree adapted from AnnoTree (Mendler et al. 2019). Inner circle represents a scale indicating the number of analyzed genomes per phylum. Black/white circles highlight abundance of genomes harboring any PSP. Most outer squares show PSP domains found across all genomes of the respective phylum. For comprehensive dataset see Table S1. **B** A phylogenetic tree applying neighbor-joining algorithm based on concatenated proteins (Parks et al. 2018) of randomly selected representatives within the order of Gammaproteobacteria is shown. Genomes were screened for PSP domain presence (Table S2, Materials and Methods). **C** Phylogenetic tree using the maximum-likelihood algorithm based on concatenated proteins (Parks et al. 2018) of genomes within the Enterobacteriaceae family is displayed. Representatives were probed for presence of PSP domains (Table S3).

The transcriptional activator PspF is a special case: while it was present in 50% of the phyla, its domain composition is not restricted to the PSP response but is instead associated with a variety of additional cellular processes (Neuwald et al. 1999). Since the PspF regulator of the PSP response has so far only been described in Enterobacteriaceae, the occurrence of orthologous proteins outside this phylogenetic group is most likely not specifically associated with the PSP response (Elderkin et al. 2002; Flores-Kim and Darwin 2016). The same accounts for the single domain protein PspE, a rhodanese that catalyzes sulfur transfer from thiosulfate to thiophilic acceptors in a variety of cellular processes beyond the involvement in PSP networks (Cheng et al. 2008; Hatahet et al. 2014). PspD and PspG, the two remaining PSP members present in the classical system of *E. coli* have the narrowest phylogenetic distribution. PspG was only found in four genomes from three phyla outside of Proteobacteria, while PspD was restricted to Proteobacteria. Since only proteobacterial genomes harbor all PSP proteins, we next focused on the PSP distribution within this phylum. First, a representative collection of 7,500 proteobacterial genomes was resolved on the taxonomic level of the order (Figure 2B, Material and Methods). Here, the complete set of PSP members was only found within Enterobacterales, whereas the remaining orders showed only partial representation of the PSP system (Figure 2B). Within the order Enterobacterales, we next resolved the PSP domain distribution at the taxonomic rank of families (Figure 2C). Remarkably, the complete PSP system, as found in *E. coli*, was only present in the closest relative species, such as *Salmonella enterica*. More distantly related species of the same family, such as *Photorhabdus luminescens*, harbors PSP associated proteins with only the core functions, comprised of PspABC. This observation already hints at the existence of alternative compositions of the PSP response system even in bacteria closely related to *E. coli* (Figure 2C) (Flores-Kim and Darwin 2016).

### PspA and PspC domains are the most prevalent PSP network members

The indicated diversity of the PSP response prompted us to next identify the PSP profiles of each genome within the dataset to establish a comprehensive overview of co-occurring PSP patterns and to resolve the conservation of PSP network organization in bacteria and archaea. Towards this goal, we analyzed the co-occurrence of PSP members in each genome. We first screened the dataset for the presence or absence of individual PSPs. 68% of the approx. 22,000 genomes encoded at least one PSP domain-containing protein, while close to 7,200 genomes lacked any PSP representative (Figure 3A and Table S1). More than 60% of the 40,000 PSP proteins identified contained either PspA or PspC domains (Figure 3A), including genomes containing multiple copies of the same PSP domain. For example, we identified eight PspA proteins in the genome of *Aneurinibacillus tyrosinisolvens*, which was recently isolated from methane-rich seafloor sediments (Tsubouchi et al. 2015) (Table S1).

**Figure 3:**
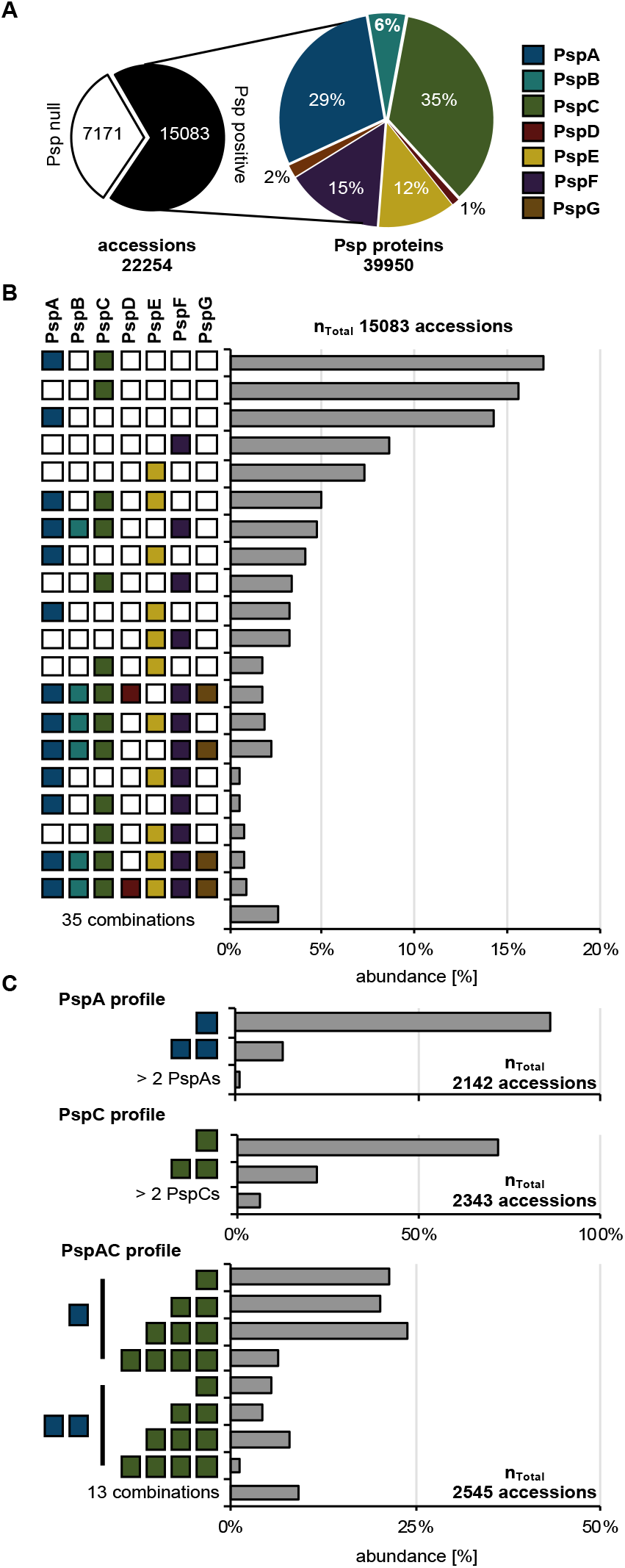
PSP profiles resolved on genomic level. **A** Screening of the full dataset for PSP domains presence/absence and their relative abundance (Table S1). **B** Descending categories of genomes according to found PSP domain profiles. Abundance was set relative to PSP positive genomes in **A** (Table S4). **C** In-depth analysis of PspA or C and PspAC containing genomes probed for abundance of multiple proteins per genome containing further PspA or C domains (Table S5).

We next categorized the PSP domain distribution and abundance by assigning a PSP profile to each genome. From approximately 15,000 genomes encoding PSP proteins, more than 45% only harbored PspA-, PspC- or PspA- and PspC-domain proteins (Figure 3B and Table S4). In comparison, only about 1% (79 genomes from Enterobacteriaceae) encoded the full repertoire of the PSP network (Figure 3B). These observations strongly support the hypothesis that most PSP networks deviate in their architecture from the proteobacterial blueprint exemplified by *E. coli* or *S. enterica*. More than 10% of the genomes containing only PspA or PspC encoded their multiple paralogs (Figure 3C and Table S5). These duplications were found in 16 phyla for PspA and 23 for PspC. In genomes harboring both PspA and PspC, many encoded multiple PspC domains. This may suggest that either the signaling properties of PspC domains serve beyond their relationship linked to PspA or that different stimuli are integrated via individual PspC proteins to enhance PSP response specificity (Figure 3C).

### Domain combinatorics and diversity of PspA and PspC

We next focused our attention on the predominant PspA- and PspC domains and domain architectures of the cognate proteins, in order to identify domain combinations that have been established and conserved in the course of evolving PSP-like responses. Such conserved domain combinations might *e.g.* provide important mechanistic insights on the regulation of PSP responses that differ from the proteobacterial model. For example in *C. glutamicum*, PspC domain is part of a histidine kinase, indicating that PspC-dependent sensing is transduced by a two-component system in order to orchestrate a PSP-like envelope stress response in this actinobacterium (Kleine et al. 2017). Towards this end, we calculated sequence lengths of PspA and PspC domain-containing proteins. Since protein domains are on average 100 AAs long, typically ranging from 50 to 200, we expected that shuffling of domain architectures within one protein would result in a notable extension of protein length (Xu and Nussinov 1998; Wheelan et al. 2000). For the PspA domain-based search, we used the Pfam PspA_IM30 HMM model of 221 AAs length (El-Gebali et al. 2019) (Material and Methods). Analysis of the length distribution of all PspA-like proteins revealed that the majority of proteins are approx. 200-250 AAs long, indicating no significant sequence space for additional domains (Figure 4A and Table S6). One notable exception was the wider size distribution of PspA homologs in the Actinobacteriota: in this phylum, numerous PspA-containing proteins were approx. 300 AA in length. However, a subsequent analysis of these proteins, using the HMMscan module, failed to identify any additional PspA-associated domains (see Table S14). In contrast, the analysis of PspC-containing proteins identified diverse domain architectures. The majority of proteins are far longer than the ~50 AAs typical of the stand-alone PspC domain (Figure 4B) (El-Gebali et al. 2019). In the Actinobacteriota, the size range of PspC-containing proteins was between 50 and 1000 AAs (Figure 4B). A detailed analysis of all protein sequences longer than 300 AAs revealed that in more than 50% of them the PspC domain was accompanied by a histidine kinase domain (see Table S7 for details), substantiating the diverse role of PspC in signal transduction processes within this phylum (Kleine et al. 2017; Ravi et al. 2018). Conserved combinations of PspC with other domains were not restricted to Actinobacteriota. In Firmicutes and Spirochaetota, we observed PspC domains arranged with several domains of unknown function (DUFs). Notably, in Firmicutes we identified proteins combining PspC and PspA domains, which demonstrates the existence of alternative PSP architectures compared to those found in *E. coli*, in which the domains are encoded by separate genes (Figure 4B and Table S7). Additionally, we identified phylum-specific (Bacteriodota and Firmicutes) C-terminal conserved regions in PcpC that did not match any protein domain model from public databases (Figure 4C and Table S8). A previous report demonstrated that the C-terminal region of PspC is of particular importance in secretin-dependent induction of a PSP response in *Y. enterocolitica* (Gueguen et al. 2009). Thus, we hypothesize that this conserved region found in representatives of Bacteriodota and Firmicutes might also perform signaling functions in their respective PSP networks. To obtain a complete picture of PSP network architectures, we next expanded our analysis to the conservation of the genomic neighborhood of PspA- and PspC-encoding genes.

**Figure 4:**
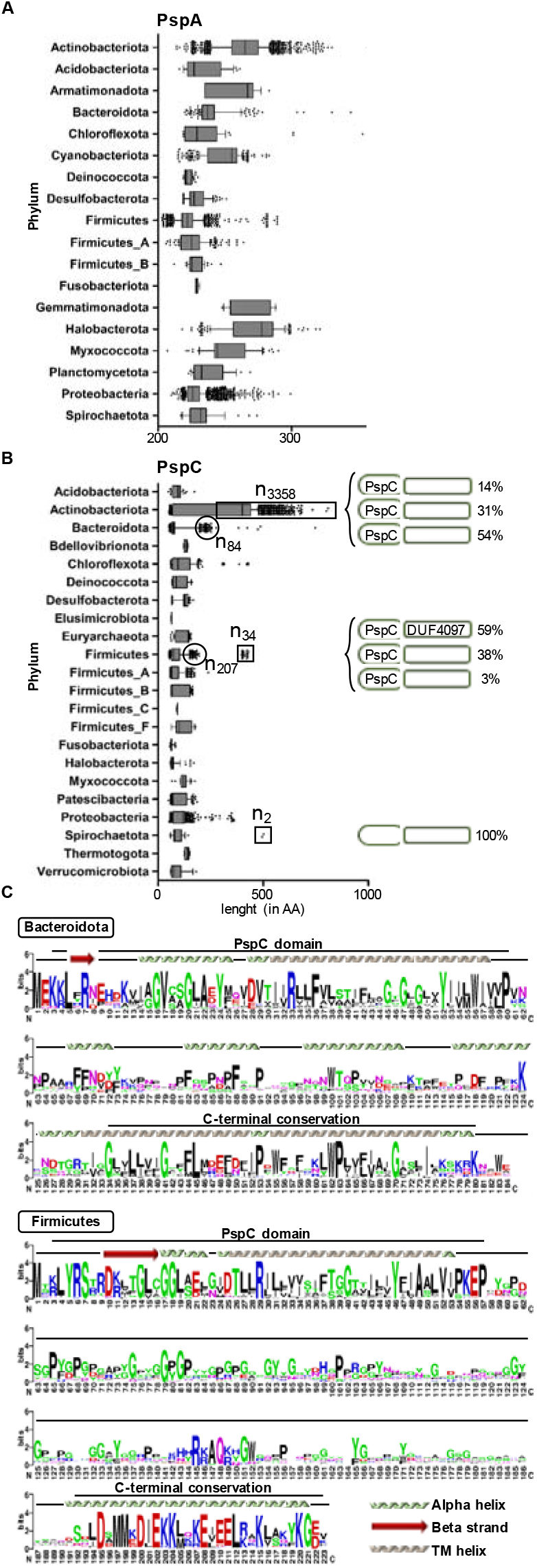
Protein length analysis of PspA and PspC. **A, B** Amino acid length distribution of PspA and C proteins found in depicted phyla. Phyla with more than ten proteins were considered (Table S6 and S7). **C** Multiple sequence alignment of C-terminal conserved regions of PspC proteins within Bacteriodota and Firmicutes. Proteins were considered as highlighted in **B** (Table S8 and File S1).

### Genomic context conservation of PspA and PspC-encoding genes

In prokaryotes, genes are often organized in operons, that encode physically interacting proteins (Koonin and Mushegian 1996; Dandekar et al. 1998; Wells et al. 2016; Esch and Merkl 2020) or proteins from the same functional pathway (Rogozin, Makarova et al. 2002, Zaslaver, Mayo et al. 2006). The systematic association between functionally related genes in operons is frequently used to characterize genes with unknown function (Overbeek et al. 1999; Wolf et al. 2001; Moreno□Hagelsieb and Janga 2008). The intergenic distance and orientation of individual genes is usually a reliable measure to predict operon arrangements (Salgado et al. 2000). It is well established that intergenic regions between identically oriented genes with less than 100-200 bp gaps enable accurate prediction of operon structures (Moreno-Hagelsieb and Collado-Vides 2002; Strong et al. 2003; Chen and Dubnau 2004; Edwards et al. 2005).

For our analyses of PspA- and PspC-encoding gene neighborhoods, an operon was defined as genes of the same orientation that are closer than 150 bp to each other (see Materials and Methods). We generated gene neighborhood profiles for phyla containing more than ten PspA or PspC proteins respectively (Figure 5A, B; for full datasets see Tables S9 and S10). Subsequently, protein sequences of the potentially co-expressed genes were retrieved, and protein domains were identified using HMMscan (see Materials and Methods). We then created consensus gene neighborhoods based on the abundance of protein domains within each phylum (Figure 5A, B and Tables S9 and S10). As expected for Proteobacteria, PspA was predominantly accompanied by PspB, PspC and PspD, reflecting the well-studied *psp* operon of *E. coli*. PspE was missing from the consensus gene neighborhood, despite its high abundance within some orders of the Proteobacteria (Figure 2B). Our analysis also reinforced previous observations that the PSP operon is often encoded with *ycjX*-like genes, containing the DUF463 domain within the phylum of Proteobacteria (Huvet et al. 2011). DUF463 belongs to the Pfam superfamily “P-loop_NTPase (CL0023)”, which contains many proteins that are involved in assembly and function of protein complexes (Neuwald et al. 1999). However, a physiological link of the PSP response with these proteins is still unknown. Moreover, DUF463-containing proteins are not mandatorily associated with the PSP response, as this domain is also found in 14% of all PSP null genomes (7171, Figure 3A), most of which belong to the phylum of Proteobacteria (see also Table S15). Beyond the Proteobacteria, the core PspABC protein set was only conserved within the phylum Desulfobacterota. In most phyla PspA is encoded without any other classical PSP protein in its neighborhood, with the exception of PspC. In a notable number of phyla, such as Acidobacteriota, Bacteriodota, Firmicutes and Fusobacteriota, proteins containing the Band 7 domain were found encoded next to *pspA* genes. The presence of *pspA* genes in actinobacterial operons encoding histidine kinases suggests alternative ways of regulating PspA-mediated (envelope) stress responses. Our analysis demonstrated an overall tendency for *pspA* genes to be co-located with genes encoding DNA binding proteins or other regulatory domain-containing proteins, *e.g.* in the phyla Acidobacteriota, Firmicutes or Spirochaetota (Figure 5A). Previously, this was observed just in a few model organisms, e.g. *pspA* is encoded adjacent to *pspF* in *E. coli* and *Y. enterocolitica* (Huvet et al. 2011). In Cyanobacteriota, more than half of the PspA proteins are encoded as mono-cistronic transcriptional units.

**Figure 5:**
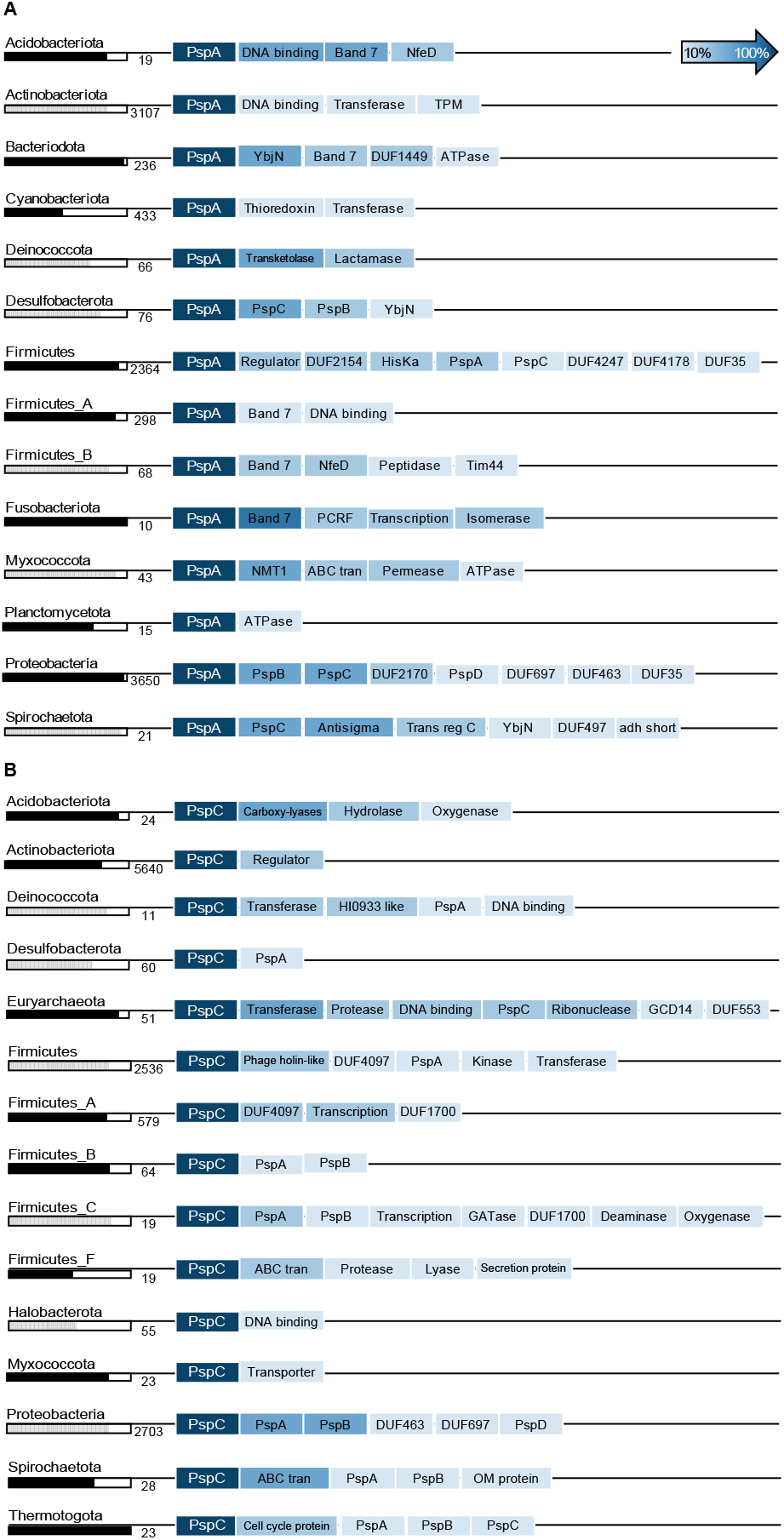
Gene neighborhood analysis of PspA and PspC. **A**, **B** Consensus gene neighborhood for PspA and C proteins based on domain abundance within depicted phyla. Black/white column indicates fraction of respective proteins encoded in operons dependent on the total amount of proteins accounted for (number next to column). For details see Material and Methods section and Tables S9 and S10.

For PspC, the gene neighborhood analysis revealed a clear preference for genes encoding predicted membrane-associated proteins (Figure 5B). In the majority of analyzed phyla, PspC was encoded together with PspA and DUF proteins as well as ABC transporters and other membrane proteins. In contrast to PspA, which is preferentially encoded in operons, PspC genes regularly locate outside of an operon structure (Figure 5B black/white bars). In phyla encoding PspC in absence of PspA, PspC-containing proteins are predominately accompanied by genes encoding transporters or DNA binding proteins. Especially in Actinobacteriota and Halobacterota, PspC appears to be primarily involved in cellular signaling without PspA contribution. This is in line with the observed presence of PspC domain as a sensory unit of histidine kinases and as part of signal transduction pathways, as discussed above (Figure 4B).

### PSP interaction network in *Bacillus subtilis*

Our analysis of a phylogenetically diverse genomic dataset revealed that PspA and PspC are the most widely distributed components of PSP response system which indicates their ancient origin. Our study demonstrated that there is no “typical” PSP network architecture: the paradigmatic PSP responses as described in *E.coli* and *Y. enterocolitica* represent just one type of the PSP system and a plethora of alternative PSP network architectures exists in the microbial world. As a first proof-of-principle, we analyzed the PSP network of the Gram-positive model organism *B. subtilis*. The genome of *B. subtilis* encodes two PspA homologs (PspA and LiaH) and one PspC homolog (YvlC), which are spread across three operons. We hypothesized that most of the additional 11 genes in these operons encode proteins that partake in the PSP-dependent CESR network (Figure 6). A tight physiological and regulatory link between the proteins encoded in the *liaIH-liaGFSR* locus has been documented (Jordan et al. 2006; Wolf et al. 2010; Dominguez-Escobar et al. 2014). While the regulatory role of the LiaFSR ‘three’-component system in orchestrating expression of the *liaIH* operon as the main effector of the Lia response is firmly established, no function could so far be attributed to LiaG. This membrane anchored protein contains a DUF4097 domain, which can also be found in another hypothetical protein, YvlB, which is encoded in the *yvlABCD* operon. Remarkably, this domain has been found in the genomic neighborhood of PspC-encoding proteins in other Firmicutes. The remaining three genes in this operon encode putative membrane proteins. The second PspA homolog, namely PspA, is encoded in the *pspA-ydjGHI* operon, where three genes encode additional membrane proteins and a cytoplasmic protein. One of the transmembrane proteins, YdjG contains a zinc ribbon domain (Pfam: TF_Zn_Ribbon) that is known to bind DNA and therefore to be involved in regulatory processes within the cell. The cytoplasmic protein YdjI belongs to the Band 7 protein family.

**Figure 6:**
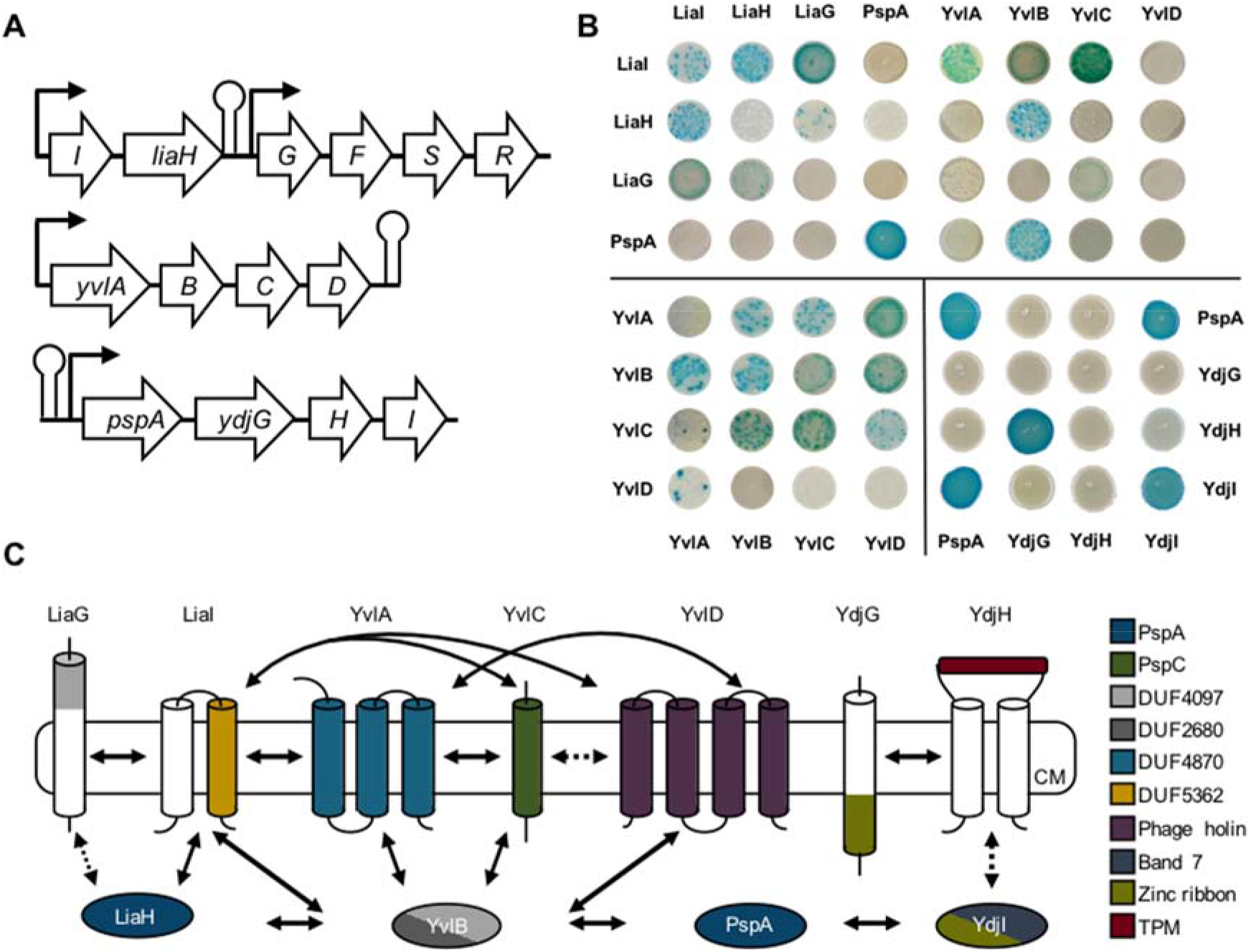
PSP network in *B. subtilis*. **A** Genomic organization of the PSP network across three separate operons in the *B. subtilis* genome. **B** Representative B2H interactions of *B. subtilis* PSP network proteins. (For full dataset see Figures S1 and S2). **C** Schematic representation of the B2H results, comprising all members of the PSP network in *B. subtilis*. Protein domains are indicated by color and protein-protein interaction highlighted as full or in case for partial interaction as dotted arrows.

For a complete picture of protein interaction network within PSP system of *B. subtilis*, we performed a bacterial-two-hybrid (B2H) screen with all proteins from three operons encoding PSP domain-containing proteins (Figures S1 and S2). The B2H is a powerful and sensitive tool for detecting not only stable interacting proteins but also weak and transient protein interactions (Stasi et al. 2015; Lin and Lai 2017). The B2H showed strong protein-protein interaction not only between proteins encoded in the same operon but also with proteins encoded in other operons (Figure 6). Membrane anchor protein LiaI strongly interacted with the PspA homolog LiaH, which substantiates a previous study demonstrating that LiaI serves as the membrane anchor for LiaH upon cell envelope damage (Wolf et al. 2010; Domínguez-Escobar et al. 2014). Moreover, we observed strong LiaI and LiaG interaction and a weak interaction of LiaG with LiaH. To date, the role of LiaG within the Lia CESR remains elusive, since a LiaG deletion has no observable phenotype (Jordan et al. 2006). Our data indicates that LiaG might promote recruitment of LiaH to the membrane either by binding the protein directly and/or supporting the LiaI-LiaH complex (Figure 6C). Furthermore, LiaG contains an extracellular DUF4097 domain containing a β-propeller motif. Beta-propeller motifs are assumed to be involved in signal transduction and protein-protein interactions (Fülöp and Jones 1999). The same domain occurs within the YvlB protein (Figure 6C). Notably, YvlB strongly interacts with PspA and LiaH as well as with other proteins encoded in the *yvl* operon (Figures 6B and 6C; Figures S1 and S2). YvlB appears to play a key role in connecting the PspA-homolog encoding operons together with the *yvl*-operon containing the PspC-homolog YvlC (Figure 6C). Regarding the protein-protein interaction between PspA and the Ydj proteins, PspA is potentially recruited to the cell membrane and seems to interact with YdjG or YdjH via the adapter protein YdjI that contains a Band 7 domain and belongs to the flotilin protein family. Flotilins are described to be involved in the organization of membrane micro domains or so called lipid rafts (Lopez and Koch 2017). In summary, our computational approach led to the identification a PSP network spanning across three different loci in *B.subtilis*.

### PSP systems in archaea

To date, PSP networks in archaea are largely unknown. We analyzed 915 archaeal genomes representing ten phyla (Table S1). Similar to bacteria, PspA and PspC proteins were both most prevalent and taxonomically widely distributed, with 25% and 12% of the analyzed genomes contained solely either protein, respectively. In contrast, more than 50% of all genomes lacked any of the PSP proteins, arguing for an overall rather low distribution of PSP members across archaea (Table S11). One archaeal genome, *Methanoperedens sp. BLZ2* classified as Halobacterota, encoded genes for two PspAs, one PspB, and one PspC protein. Strikingly, all these proteins are encoded in separate operons. PspA proteins are located next to genes encoding a small multi-drug exporter or an extracellular peptidase. PspB and PspC are parts of operons that also contain genes encoding for multiple DNA-binding HTH domains and combined PAS domain-containing or response regulator proteins indicating their involvement in signal transduction. Proteomic analysis of *Haloferax volcanii* revealed upregulation of a PspA homolog upon salinity stress (Bidle et al. 2008). This observation indicates a functional overlap between PSP responses in archaea and bacteria. Applying our thresholds for an operon structure (see Materials and Methods), we identified the fourth transmembrane protein encoded downstream of *pspA* (locus tag: Hvo2637). Subsequent analysis identified two domains in this protein, bPH_4 and EphA2_TM, with probabilities of 60% and 80%, respectively (Adebali et al. 2015). EphA2_TM has been associated with tyrosine kinase acceptors (Bocharov et al. 2010), which suggests that the protein might potentially play a signaling role within the PSP response of *H. volcanii*. PSP members seem poorly conserved within archaeal representatives. Most genomes do not encode PSP associated genes, whereas the majority of those that encode are limited to PspA or PspC.

### Concluding Remarks

Previous studies on the diversity of the PSP system were limited to 3 bacterial phyla (Huvet et al. 2011; Manganelli and Gennaro 2017; Ravi et al. 2018). We substantially expanded previous analyses by carrying out an unbiased comparative genomic analysis of PSP response networks across more than 100 phyla of bacteria and archaea. We performed PSP domain model-based searches and revealed a variety of PSP architectures and a heterogeneous distribution of PSP proteins across taxonomic groups. We demonstrated that PspA and PspC are both the most frequently found and phylogenetically most widely distributed components of the PSP response network (Figure 2A). Our analysis revealed diverse protein networks of PSP responses across bacteria and archaea. The PSP system displays remarkable diversity with respect to component design, phylum-specific conserved architectures, and postulated modes of both signal transduction and underlying physiologies. We fully reconstructed *in silico* PSP networks in two phyla that contain large number of sequenced genomes – Acidobacteriota and Bacteroidota. For example, in Acidobacteriota, we found PspA associated with Band 7 and FloT domain containing proteins responsible for stabilizing membrane integrity upon changes of its fluidity state under stress conditions (Bach and Bramkamp 2013). The NfeD protein, which has been identified as partner-protein in gene neighborhood analyses, may support PspA assembly and/or function (Green et al. 2009). We also identified the HTH domain containing transcriptional regulators encoded in the PspA operons that potentially can regulate the operon expression. Our analysis suggests that PspC in Bacteroidota might be directly involved in the beta-lactam stress response (Figure 7).

**Figure 7:**
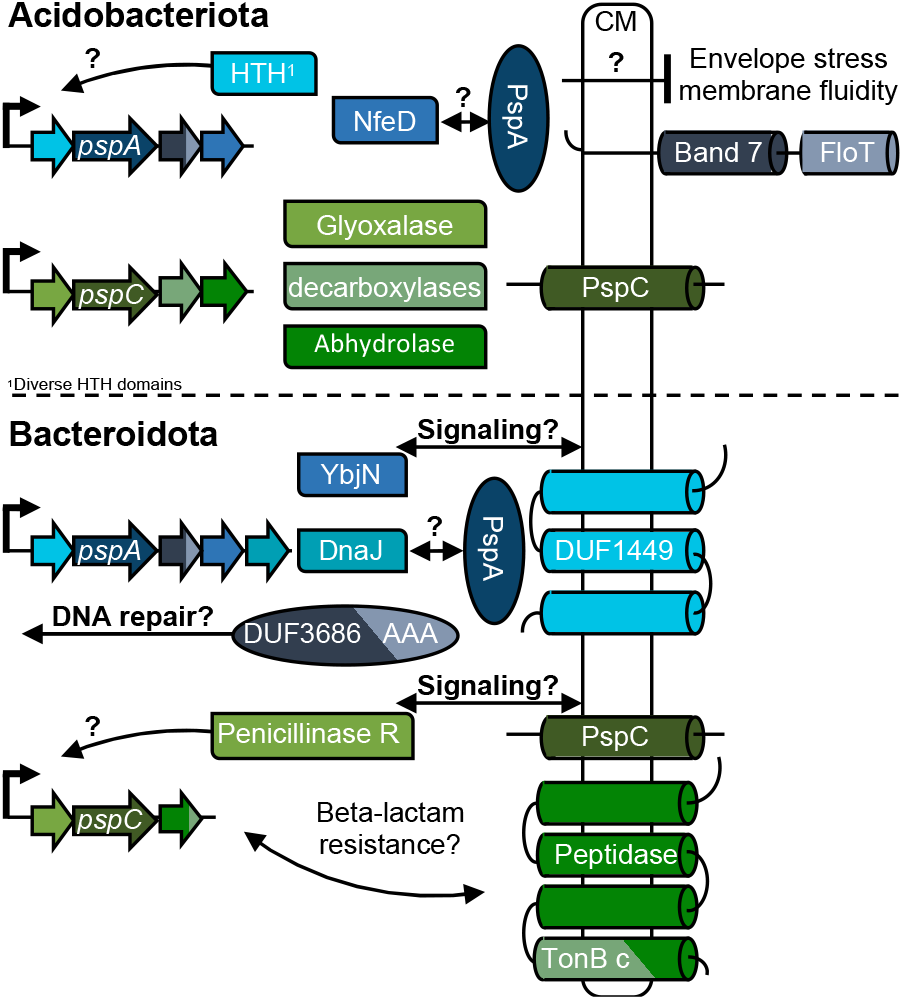
PSP network predictions. Phylum specific *in silico* predictions of PspA and PspC networks. Potential protein-protein interactions are indicated by arrows and physiological roles are implied. Data derives from gene neighborhood, HMM scan and THMM analyses, for full dataset see Table S12.

We demonstrated the utility of our computational analysis by showing that proteins predicted to comprise the PSP system in *B. subtilis* physically interact, although they are encoded in three different operons (Figure 6 and Figures S1 and S2). It is well known that the Lia system containing the PspA homolog LiaH serves as a resistance determinant under CES conditions (Mascher et al. 2003; Mascher et al. 2004; Radeck et al. 2016). Interaction of LiaH with the YvlB protein revealed in this study suggests a potential additional role for LiaH within the PSP network.

In summary, our comprehensive and unbiased analysis of bacterial and archaeal genomes revealed an unexpected diversity of the PSP system that can now be tested experimentally.

## Materials and Methods

### Bioinformatics tools and software environments

The following software packages were applied in this study: HMMER v3.2.1 (Eddy 2011), MAFFT 7 online version (Katoh et al. 2017), MEGA-X (Kumar et al. 2018), CDvist (Adebali et al. 2015), Jalview (Waterhouse et al. 2009), iTOL (Letunic and Bork 2019) and CD-HIT (Huang et al. 2010). Multiple-sequence alignments were generated using the default set L-IN-I algorithm from MAFFT (Katoh et al. 2017). The neighbor-joining tree (Figure 2B; Supplement table 2) was built in MEGA-X with pairwise deletion using the poisson model. All maximum likelihood trees were computed in MEGA-X with applying the Jones-Taylor-Thornton (JTT) substitution model and using all sites, unless otherwise specified. Final visualization and mapping of PSP domains onto the phylogenetic trees was performed using iTOL. Computational analyses were executed on a local computing environment and custom scripts for data processing, filtering and evaluation were written in Python v2.7 or v3.7 in the environment of PyCharm v2019.1.1.

### Data sources

Bacterial and archaeal genomes were analyzed that are declared as representatives according to the Genome Taxonomy Database (GTDB) r86 (Parks et al. 2018). In total, 22,254 proteomes were obtained that were available at the National Center for Biotechnology Information (NCBI) Reference Sequence (RefSeq) or GenBank databases as of October 2018 (Supplement table 1). All performed analyses were done using this final dataset or derived subsets as indicated. Analysis of PSP domains in the obtained genomes was performed using HMMer (http://hmmer.org/). To run the HMMsearch module, hidden-markov-models (HMM) were obtained from the Protein Families database (Pfam): PF04012.12, PF06667.12, PF04024.12, PF09584.10 and PF09583.10 (El-Gebali et al. 2019). Models for the two remaining PSP related proteins, PspE and PspF, for which no model was available at Pfam, were generated by downloading their TIGRFAMs, TIGR02981 and TIGR02974 respectively (Haft et al. 2001). The seed sequences were obtained and used to build HMM models locally using the HMMbuild module.

### Identification of PSPs in the genomic datasets

All seven, PspA to PspG, HMM models were probed against each genome downloaded applying the HMM- search module. Obtained results were checked for their accuracy and to avoid false positive hits via a python script that filtered following criteria: i) the e-value threshold cutoff for a positive hit was set to 10^−3^, ii) the domain had to start within the first 50 AA of the protein and iii) the alignment of the HMM model with the protein sequence had to exceed 90% of the models length. In total 39,950 proteins fulfilled these criteria and were considered as PSPs (Supplement Table 1).

### Construction of phylogenetic trees

The sets of concatenated and aligned 120 proteins (bacterial) and 122 (archaeal), proposed by the GTDB criteria, were used to build phylogenetic trees in Figure 2 (Parks et al. 2018). Figure 2A was adapted and modified from the pre-computed phylogenetic tree presented by AnnoTree (Mendler et al. 2019) and only serves as representation of the phylogenetic distribution of PSPs. For the dataset in Figure 2B, a maximum of 55 members of each of the 53 orders within the class of Gammaproteobacteria were randomly chosen resulting in 742 genomes (Supplement table 2). The amount of 55 genomes was set according to a hyper geometric distribution setting a 95% probability threshold to at least include one genome within the Enterobacterales order having the full set of PSPs. From orders with less than 55 members all genomes were included and a neighbor-joining tree was computed. The dataset in Figure 2C, is based on the species (41 genomes) used in (Ortega and Zhulin 2016), representing the diversity of the Enterobacteriaceae family (Supplement table 3). Four species had to be replaced by genomes declared as representatives according to the GTDB. A maximum likelihood tree using the JTT model with 100 bootstraps was applied to the respective set of 120 concatenated aligned proteins provided by the GTDB.

### Protein domain co-occurrence analysis

To characterize potential co-occurring protein domains associated within PspA or PspC domain-containing proteins, respective proteins were downloaded at the NCBI protein database (Supplement Table 6 and 7). The HMMscan module was used with an e-value threshold of 0.001 to search for associated domains in PspA and PspC domain-containing proteins. Identified domains were obtained from the Pfam HMM (march 2019) library for PFam-A families. For better overview, only phyla containing more than ten PSPs were included into the Figures 3A and B respectively. For full dataset see Supplement Table 6 and 7.

### Creation of Weblogos for PspC domain-containing proteins

To identify PspC C-terminal conserved regions, respective proteins were further analyzed by generating a multiple sequence alignment (MSA) in MAFFT (Katoh et al. 2017) and Jalview for visualization (Waterhouse et al. 2009) (Figure 4C). The weblogo of the final MSA was created using the online version of Weblogo (Crooks et al. 2004). Secondary structure predictions were performed in Phyre2 (Kelley et al. 2015). For used proteins, see Supplement Table 8 and the MSA provided in Supplement File S1.

### Gene neighborhood analysis

Analysis of gene neighborhood was performed using the application programming interface (API) implemented in the Microbial Signal Transduction database (MiST3) (Gumerov et al. 2020). For each query gene, five up- and downstream genes were obtained and filtered for orientation and their respective location. An operon structure was defined for genes that shared the same strand orientation and were located no more than 150 base pairs apart from each other. Proteins that were identified as gene neighborhood were then obtained from the Refseq NCBI database. The HMMscan module was then used to identify protein domains, using an e-value threshold of 0.001. To exclude overlapping domain hits, e.g. Band_7 and Band_7_1, which would cover the same sequence space, only first domain hits were considered. For proteins, containing multiple non-overlapping domains, their domain architecture was fused *e.g.* HisKA_3 and HATPase_c and finally categorized (Supplement Tables 9 and 10). For better overview and to generate the consensus gene neighborhood, two thresholds were applied: i) only phyla containing at least ten PSP positive genomes were included and ii) the identified protein domain had to be present in more than 10% of the genomes within the analyzed phyla *e.g.* from the 19 PspAs analyzed in Acidobacteriota, 11 contained a Band 7_Flot protein in their gene neighborhood, thus resulting in 58%. For complete dataset see Supplement Tables 9 and 10.

### DNA manipulation

Plasmids were constructed using standard cloning techniques as described elsewhere (Sambrook and Russell 2001). For DNA amplification via PCR, Q5 polymerase was used. Enzymes were purchased from New England Biolabs (NEB, Ipswich, MA, USA) and applied following their respective protocols. Positive *E. coli* clones were checked by colony PCR, using OneTaq polymerase. All constructs were verified by sequencing. All strains, primers and plasmids used in this chapter are listed in Supplement Table 13.

### Bacterial-two-hybrid assay

The bacterial-two-hybrid experiment is based on an adenylate cyclase reconstruction resulting in the transcription of the reporter gene *lacZ* in *E. coli* (Karimova et al. 1998). The gene is encoding for a β-galactosidase. The enzyme is able to cleave X-Gal that results in blue colored colonies. For the bacterial-two-hybrid-assay, the adenylate cyclase is divided into two parts, each of them either N- or C-terminal present on the vectors pUT18/pUT18C or pKT25/pKT25N. Genes of interest, encoding candidates for protein-protein interaction, were cloned into these vectors and a co-transformed into *E. coli* BTH101. Because the interaction of the proteins can be influenced by the position of the adenylate cyclase, all plasmid combinations were used. The vectors pUT18 zip and pKT25N zip were included and served as a positive control and the empty vectors pUT18 and pKT25 were used as a negative control. After the transformation, the cells were pelleted and resuspended in 40 μl LB medium. 10 μl of each transformation mix was spotted on agar plates containing Ampicillin (100 μg ml^−1^), Kanamycin (50 μg ml^−1^), IPTG (0.5 mM) and X-Gal (40 μg ml^−1^). After the spots dried, the procedure was repeated. The plates were incubated at 30°C overnight and the next day the rest of the transformation mix was spotted two times. In most of the cases not enough colonies grew to cover the whole spot area. To achieve a higher colony density, overnight cultures of the different strains were prepared and 2 × 10 μl were spotted the following day. The plates were wrapped in aluminum foil to protect them from incident light exposure and stored in the fridge for several weeks to increase contrast and intensity of the colony color.

## Data Availability Statement

The data underlying this article are available in the article, in its online supplementary material and on request to the corresponding authors.

## Author contributions

P.F.P. and T.M. conceptualized the study; P.F.P. performed the bioinformatics research; D.W. and L.B. performed the bacterial two hybrid assay; P.F.P., V.M.G, E.P.A., L.B., T.M. I.B.Z., and D.W. analyzed data; P.F.P., T.M., I.B.Z., and D.W. wrote the paper with contribution from V.M.G. and E.P.A.

## Acknowledgments

The authors thank Josue Flores-Kim for personal communication on *Y. enterocolitica* PSP response. We also thank Marc Bramkamp (University of Kiel, Germany) and Robyn Emmins (University of Newcastle, UK) for plasmids as well as Elisa Granato and Mona Steichele (LMU Munich, Germany) for technical support.

