## Supplemental Figures for "Phyletic distribution and diversification of the Phage Shock Protein stress response system in bacteria and archaea"

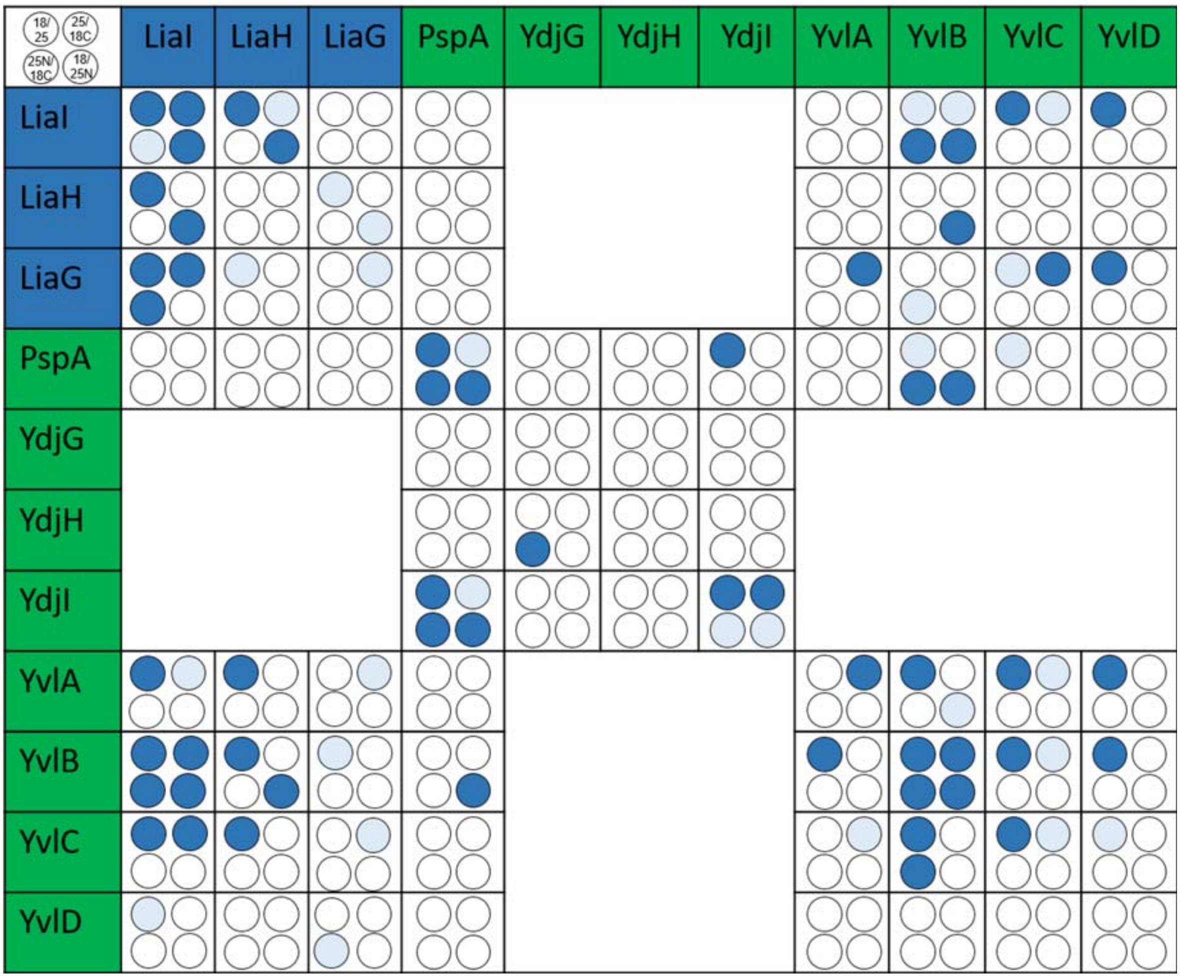


Figure S1

Overview and Summary of protein-protein interactions within the PSP network in *Bacillus subtilis* determined by B2H assays. Protein encoding genes of the Lia-system regulated by the LiaRS two-component system are marked in blue. Protein expression of loci containing the *yvl*- or *pspA*-*ydj* genes that are the control of the extracytoplasmic function factor $\sigma^W$, highlighted in green. Dark blue filled circles indicate strong protein-protein interaction whereas circles in light blue point weak interactions. White circles stand for no protein-protein interaction. For experimental details please see the Material and Methods section.

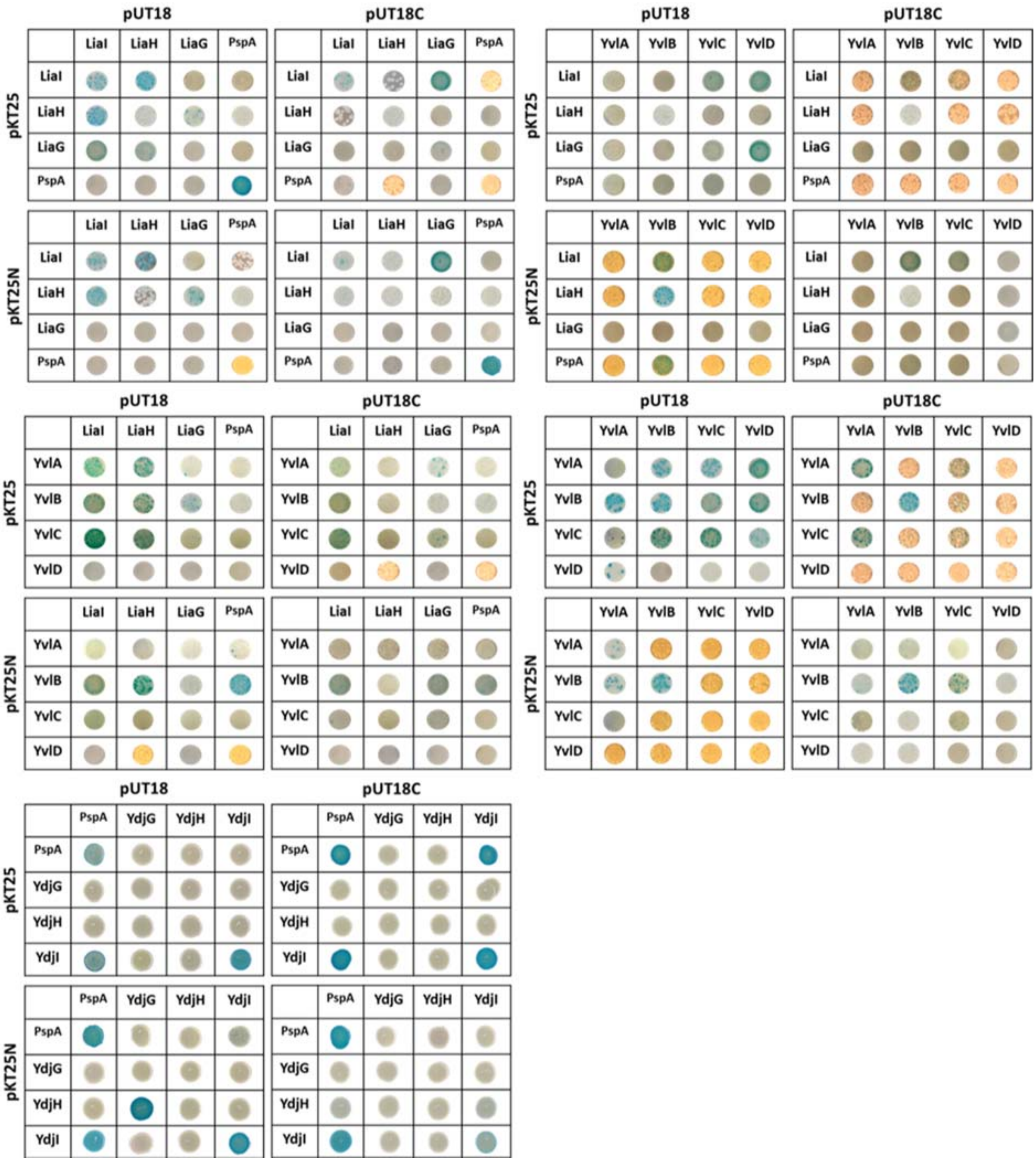


Figure S2

Colony colour of protein combinations within the PSP response in *Bacillus subtilis* tested by B2H assay. Please see Material and Methods for experimental details.
