## Supplemental File_S1 for "Phyletic distribution and diversification of the Phage Shock Protein stress response system in bacteria and archaea"

Phyre2

| Email | |
|---|---|
| Description | Weblogo_Bacteriodota |
| Date | Wed Sep 11 15:40:48 BST 2019 |
| Unique Job ID | 7cf38cdd617c6372 |

#### Secondary structure and disorder prediction

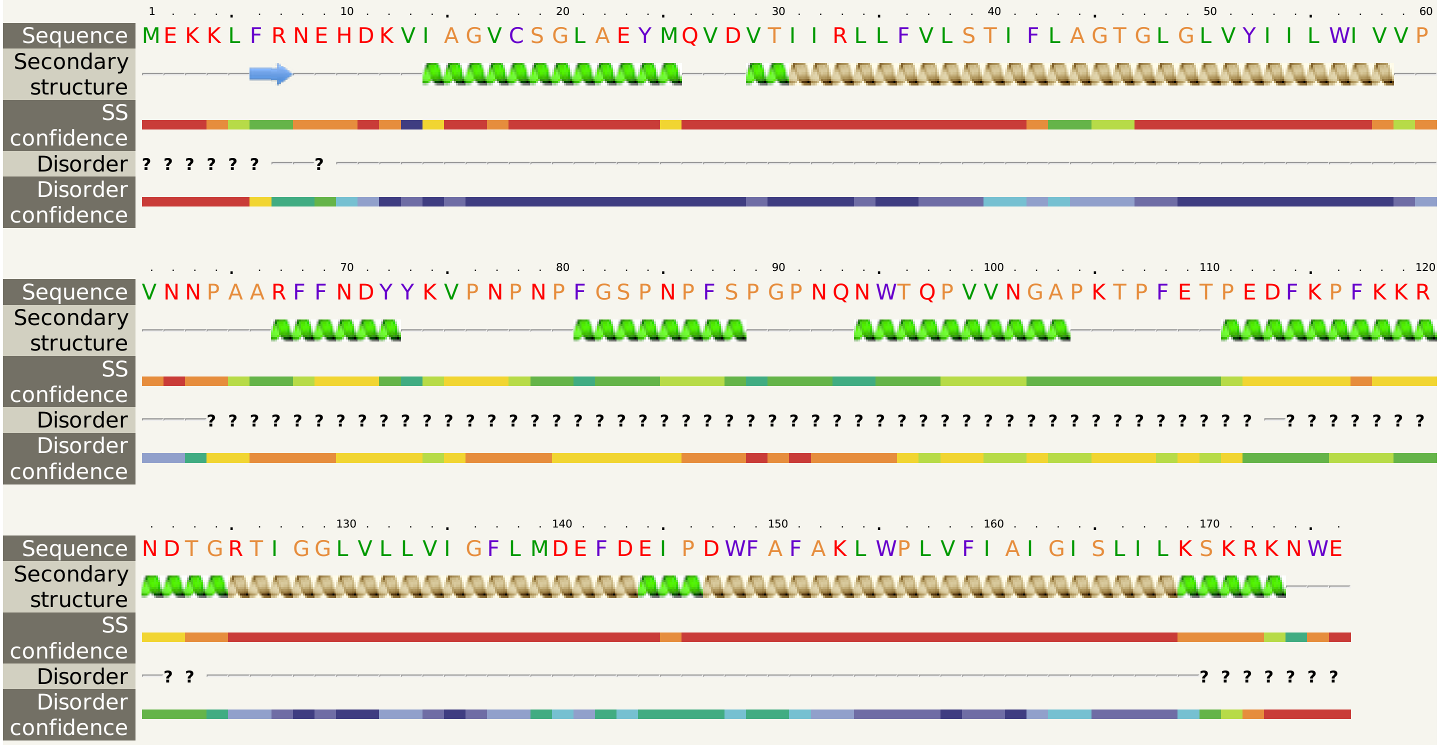


Confidence Key

High(9) Low (0)

? Disordered ( 41%)

Alpha helix ( 73%)

Beta strand ( 1%)

TM helix ( 39%)

### Phyre2

| Email | |
|---|---|
| Description | Weblogo_FIRMICUTES |
| Date | Wed Sep 11 15:44:30 BST 2019 |
| Unique Job ID | a1a798ec058c7191 |

#### Secondary structure and disorder prediction

Sequence (1–60): MTKLYRSTRDKLTGLCGGLAELGIDTLLRILLVVSIFTGGTTILIYFIAALVIPKEPYGP

Disorder: ? ? ? ? ? ? ? (1–7); ? ? ? ? ? (56–60)

Sequence (61–120): DSGPYGPGPGAPYGPYGGPGPYYGPGPGGYYGYDHGPPRGPYNNGGYGDPGGGGYGPPGG

Disorder: ? (61–120, all residues)

Sequence (121–180): GAYGGRPPKHHRAQKHGWGPPQPGGYYYGPYGAGGAAAADSDLDSMMKDIEKKLQKEIEE

Disorder: ? (121–159)

Sequence (181–192): LRAKLAKYKGEV

Disorder: ? ? ? ? ? (188–192)

Confidence Key

High(9) … Low (0)

? Disordered ( 60%)

Alpha helix ( 34%)

Beta strand ( 3%)

TM helix ( 15%)
