## Supplemental Table_S13 for "Phyletic distribution and diversification of the Phage Shock Protein stress response system in bacteria and archaea"

### Tables

Table 1. Strains used in this study

| **Strain** | **Genotype** [a] | **Source** |
|---|---|---|
| ***Escherichia coli*** | | |
| BTH 101 | F$^-$ *cya*-99 *araD*139 *galE*15 *galK*16 *rpsL*1 (*str*$^R$) *hsdR*2 *mcrA*1 *mcrB*1 | Lab stock |
| DH10β | F$^-$ *mcr*A Δ(*mrr*-*hsd*RMS-*mcr*BC) φ80*lac*ZΔM15 Δ*lac*X74 *rec*A1 *end*A1 *ara*D139 Δ(*ara-leu*)7697 *gal*U *gal*K λ$^-$ *rps*L(*str*$^R$) *nup*G | Lab stock |
| XL1 blue | *rec*A1 *end*A1 *gyr*A96 *thi*-1 *hsd*R17 *sup*E44 *rel*A1 lac [F$'$ *pro*AB *lac*I$^q$ ZΔM15 Tn10 (*tet*$^R$) | Lab stock |
| TME183 | XL1 blue pUT18-*liaI* | This study |
| TME184 | XL1 blue pUT18C-*liaI* | This study |
| TME185 | XL1 blue pKT25-*liaI* | This study |
| TME186 | XL1 blue pKT25N-*liaI* | This study |
| TME203 | XL1 blue pUT18-*liaG* | This study |
| TME204 | XL1 blue pUT18C-*liaG* | This study |
| TME205 | XL1 blue pKT25-*liaG* | This study |
| TME206 | XL1 blue pKT25N-*liaG* | This study |
| TME207 | XL1 blue pUT18-*liaH* | This study |
| TME208 | XL1 blue pUT18C-*liaH* | This study |
| TME209 | XL1 blue pKT25-*liaH* | This study |
| TME210 | XL1 blue pKT25N-*liaH* | This study |
| TME343 | XL1 blue pUT18-*pspA* | This study |
| TME344 | XL1 blue pUT18C-*pspA* | This study |
| TME345 | XL1 blue pKT25-*pspA* | This study |
| TME346 | XL1 blue pKT25N-*pspA* | This study |
| TME554 | XL1 blue pKT25N-*yvlA* | This study |
| TME555 | XL1 blue pKT25-*yvlA* | This study |
| TME556 | XL1 blue pKT25N-*yvlB* | This study |
| TME557 | XL1 blue pKT25-*yvlB* | This study |
| TME558 | XL1 blue pKT25N-*yvlC* | This study |
| TME559 | XL1 blue pKT25-*yvlC* | This study |
| TME560 | XL1 blue pKT25N-*yvlD* | This study |
| TME561 | XL1 blue pKT25-*yvlD* | This study |
| TME566 | XL1 blue pUT18-*yvlA* | This study |
| TME567 | XL1 blue pUT18C-*yvlA* | This study |

| | | |
|---|---|---|
| TME568 | XL1 blue pUT18-*yvlB* | This study |
| TME569 | XL1 blue pUT18C-*yvlB* | This study |
| TME570 | XL1 blue pUT18-*yvlC* | This study |
| TME571 | XL1 blue pUT18C-*yvlC* | This study |
| TME572 | XL1 blue pUT18-*yvlD* | This study |
| TME573 | XL1 blue pUT18C-*yvlD* | This study |
| TME2876 | DH10β pUT18C-*ydjG* | This study |
| TME2877 | DH10β pUT18C-*ydjH* | This study |
| TME2878 | DH10β pUT18C-*ydjI* | This study |
| TME2880 | DH10β pUT18-*ydjG* | This study |
| TME2881 | DH10β pUT18-*ydjH* | This study |
| TME2882 | DH10β pUT18-*ydjI* | This study |
| TME2884 | DH10β pKT25-*ydjG* | This study |
| TME2885 | DH10β pKT25-*ydjH* | This study |
| TME2886 | DH10β pKT25-*ydjI* | This study |
| TME2887 | DH10β pKT25N-*ydjG* | This study |
| TME2888 | DH10β pKT25N-*ydjH* | This study |
| TME2889 | DH10β pKT25N-*ydjI* | This study |

[a] str streptomycin; tet tetracycline

Table 2. Plasmids used in this study

| **Name/Number** | **Genotype** [a] | **Primer** | **Source** |
|---|---|---|---|
| pUT18 | P*lac*, MCS, T18 (AA 225 to 399 of ccaA), *bla*, Ori ColE1 | -- | Karimova et al., 1998 |
| pUT18C | P*lac*, T18 (AA 225 to 399 of ccaA), MCS, *bla*, Ori ColE1 | -- | Karimova et al., 1998 |
| pKT25 | P*lac*, T25 (first 224 AA of ccaA), MCS, *kan*, Ori p15A | -- | Karimova et al., 1998 |
| pKT25N | P*lac*, MCS T25 (first 224 AA of ccaA) *kan*, Ori p15A | -- | Karimova et al., 1998 |
| pUT18 zip | P*lac*, T18 (AA 225 to 399 of ccaA)-leucine zipper of GCN4, MCS, *bla*, Ori ColE1 | -- | Karimova et al., 1998 |
| pUT25 zip | P*lac*, T25 (first 224 AA of ccaA)-leucine zipper of GCN4, MCS, *kan*, Ori p15A | -- | Karimova et al., 1998 |
| 248 | pUT18-*liaI* | -- | R. Emmins, Newcastle |
| 249 | pUT18C-*liaI* | -- | R. Emmins, Newcastle |
| 250 | pKT25-*liaI* | -- | R. Emmins, Newcastle |

| | | | |
|---|---|---|---|
| 251 | pKT25N-*liaI* | -- | R. Emmins, Newcastle |
| 272 | pUT18-*liaG* | -- | R. Emmins, Newcastle |
| 273 | pUT18C-*liaG* | -- | R. Emmins, Newcastle |
| 274 | pKT25-*liaG* | -- | R. Emmins, Newcastle |
| 275 | pKT25N-*liaG* | -- | R. Emmins, Newcastle |
| 310 | pKT25N-*liaH* | TM12337TM1234 | This study |
| 311 | pKT25-*liaH* | TM12337TM1234 | This study |
| 312 | pUT18C-*liaH* | TM12337TM1234 | This study |
| 313 | pUT18-*liaH* | TM12337TM1234 | This study |
| 543 | pUT18-*yvlA* | TM1924/TM1925 | This study |
| 544 | pUT18C-*yvlA* | TM1924/TM1925 | This study |
| 545 | pKT25N-*yvlA* | TM1924/TM1925 | This study |
| 546 | pKT25-*yvlA* | TM1924/TM1925 | This study |
| 547 | pUT18-*yvlB* | TM1926/TM1927 | This study |
| 548 | pUT18C-*yvlB* | TM1926/TM1927 | This study |
| 549 | pKT25-*yvlB* | TM1926/TM1927 | This study |
| 550 | pKT25N-*yvlB* | TM1926/TM1927 | This study |
| 551 | pUT18-*yvlC* | TM1928/TM1929 | This study |
| 552 | pUT18C-*yvlC* | TM1928/TM1929 | This study |
| 553 | pKT25-*yvlC* | TM1928/TM1929 | This study |
| 554 | pKT25N-*yvlC* | TM1928/TM1929 | This study |
| 555 | pUT18-*yvlD* | TM1930/TM1931 | This study |
| 556 | pUT18C-*yvlD* | TM1930/TM1931 | This study |
| 557 | pKT25-*yvlD* | TM1930/TM1931 | This study |
| 558 | pKT25N-*yvlD* | TM1930/TM1931 | This study |
| 769 | pUT18-*pspA* | TM1323/TM1324 | This study |
| 770 | pUT18C-*pspA* | TM1323/TM1324 | This study |
| 771 | pKT25-*pspA* | TM1323/TM1324 | This study |
| 772 | pKT25N-*pspA* | TM1323/TM1324 | This study |
| 2485 | pUT18C-*ydjH* | -- | M. Bramkamp, Kiel |
| 2486 | pUT18C-*ydjI* | -- | M. Bramkamp, Kiel |
| 2488 | pUT18-*ydjG* | -- | M. Bramkamp, Kiel |
| 2489 | pUT18-*ydjH* | -- | M. Bramkamp, Kiel |
| 2490 | pUT18-*ydjI* | -- | M. Bramkamp, Kiel |
| 2492 | pKT25-*ydjG* | -- | M. Bramkamp, Kiel |
| 2493 | pKT25-*ydjH* | -- | M. Bramkamp, Kiel |

| | | | |
|---|---|---|---|
| 2494 | pKT25-*ydjI* | -- | M. Bramkamp, Kiel |
| 2496 | pKT25N-*ydjG* | -- | M. Bramkamp, Kiel |
| 2497 | pKT25N-*ydjH* | -- | M. Bramkamp, Kiel |
| 2498 | pKT25N-*ydjI* | -- | M. Bramkamp, Kiel |

Table 3. Oligonucleotides used in this story

| **Name** | **Sequence**[a] |
|---|---|
| **Bacterial Two Hybrid Cloning** | |
| TM1233 (*liaH* fwd (*Xba*I)) | AGCG**TCTAGA**GATGGTATTAAAAAGAATCAG |
| TM1234 (*liaH* rev (*Bam*HI)) | AGCT**GGATCC**AGTTCATTTGCCGCTTTTGTCTGGTC |
| TM1323 (*pspA* fwd (*Xba*I)) | AGCT**TCTAGA**GATGAGTATAATTGGAAG |
| TM1324 (*pspA* rev (*Bam*HI)) | AGCT**GGATCC**CTCTTATCGAGCATCATTTTCGC |
| TM1924 (*yvlA* fwd (*Xba*I)) | AGCT**TCTAGA**CTTGAACCGTAATCAAGC |
| TM1925 (*yvlA* rev (*Bam*HI)) | AGCT**GGATCC**GATGCTGCACGCAGGACCTTAATG |
| TM1926 (*yvlB* fwd (*Xba*I)) | AGCT**TCTAGA**CATGAAGCAAGAAAAGGAACGAATCC |
| TM1927 (*yvlB* rev (*Bam*HI)) | AGCT**GGATCC**AACCTCTGTGAGTACTTTAG |
| TM1928 (*yvlC* fwd (*Xba*I)) | AGCT**TCTAGA**CATGAATAAGCTTTATCGCTCAG |
| TM1929 (*yvlC* rev (*Bam*HI)) | AGCT**GGATCC**AGTTTCATATCCCTTTCTGACGG |
| TM1930 (*yvlD* fwd (*Xba*I)) | AGCT**TCTAGA**CATGGTAAAATGGGCAGTCAGC |
| TM1931 (*yvlD* rev (*Bam*HI)) | AGCT**GGATCC**GGTTTTTTTCTAAGCGGCTCTAAAATGCC |
| **Bacterial Two Hybrid Cloning Check** | |
| TM1220 (pUT18 fwd) | AGCTCACTCATTAGGCACCC |
| TM1221 (pUT18 rev) | CCGTCGTAGCGGAACTGGCG |
| TM1222 (pUT18C fwd) | TCGACGATGGGCTGGGAGCC |
| TM1223 (pUT18C rev) | AGCAGACAAGCCCGTCAGGG |
| TM1224 (pKT25 fwd) | GGCGGATATCGACATGTTCG |
| TM1225 (pKT25 rev) | ATCGGTGCGGGCCTCTTCGC |
| TM1226 (pKT25N fwd) | GCTCACTCATTAGGCACCCC |
| TM1227 (pKT25N rev) | GGCGGAACATCAATGTGGCG |

[a] **Bold** = restriction enzyme recognition sites
